# Epigenetics biomarkers of delirium: immune response, inflammatory response and cholinergic synaptic involvement evidenced by genome-wide DNA methylation analysis of delirious inpatients

**DOI:** 10.1101/729335

**Authors:** Taku Saito, Hiroyuki Toda, Gabrielle N. Duncan, Sydney S. Jellison, Tong Yu, Mason J. Klisares, Sophia Daniel, Allison Andreasen, Lydia Leyden, Mandy Hellman, Eri Shinozaki, Sangil Lee, Aihide Yoshino, Hyunkeun R. Cho, Gen Shinozaki

**Affiliations:** Department of Psychiatry, University of Iowa, Iowa City, Iowa, USA; Department of Psychiatry, School of Medicine, National Defense Medical College, Tokorozawa, Saitama, Japan; Department of Internal Medicine, University of Iowa, Iowa City, Iowa, USA; Department of Emergency Medicine, University of Iowa, Iowa City, Iowa, USA; Department of Neurosurgery, Carver College of Medicine, University of Iowa, Iowa City, Iowa, USA; Department of Biostatistics, College of Public Health, University of Iowa, Iowa City, Iowa, USA; Iowa Neuroscience Institute, University of Iowa, Iowa City, Iowa, USA; Interdisciplinary Graduate Program in Neuroscience, University of Iowa, Iowa City, Iowa, USA

**Keywords:** delirium, epigenetics, genome-wide DNA methylation, immune response, inflammatory response, cholinergic synapse

## Abstract

**Background:** The authors previously hypothesized the role of epigenetics in pathophysiology of delirium, and tested DNA methylation (DNAm) change among pro-inflammatory cytokines along with aging in blood, glia and neuron. The authors reported that DNAm level of the *TNF-alpha* decreases along with aging in blood and glia, but not in neuron; however, DNAm differences between delirium cases and non-delirium controls have not been investigated directly. Therefore, in the present study, DNAm differences in blood between delirium patients and controls without delirium were examined.

**Methods:** A case-control study with 92 subjects was conducted. Whole blood samples were collected and genome-wide DNAm was measured by the Infinium HumanMethylationEPIC BeadChip arrays. The correlation between DNAm levels in the *TNF-alpha* and age, network analysis, and the correlation between age and DNAm age were tested.

**Results:** Only delirium cases showed 3 CpGs sites in the *TNF-alpha* significantly correlated to age after multiple corrections. A genome-wide significant CpG site near the gene of *LDLRAD4* was identified. In addition, network analysis showed several significant pathways with false discovery rate adjusted *p*-value < 0.05. The top pathway with GO was immune response, and the second top pathway with KEGG was cholinergic synapse. Although there was no statistically significant difference, DNAm age among non-delirium controls showed “slower aging” compared to delirium cases.

**Conclusions:** DNAm differences were shown both at gene and network levels between delirium cases and non-delirium controls. This finding indicates that DNAm status in blood has a potential to be used as epigenetic biomarkers for delirium.

## INTRODUCTION

Delirium among elderly patients is dangerous and common—it occurs in 15–53% of elderly patients after surgery, and in 70–87% of those in intensive care (1); however, it is underdiagnosed and undertreated (2). Delirium is notorious for its association with a long term cognitive decline (3) and high mortality (4, 5). Given that our society is aging, it is becoming increasingly important to predict which patients are at risk of experiencing delirium. Previous searches for biomarkers have revealed increased serum levels of inflammatory markers (6, 7) and cytokines (8, 9) among elderly patients with delirium. Similarly, studies in animal models have shown that cognitive disturbances (10) in response to exogenous insults increase with age and that cytokine release from microglia play a key role (11). Thus, it is possible that microglia and inflammatory cytokines play a role in the pathogenesis of delirium in humans. However, the mechanisms whereby cytokine release or neuro-inflammation is enhanced with aging remain unclear, and identifying the patients in whom this occurs and optimizing their care will require reliable biomarkers.

Given the fact that aging and inflammation are the key risk factors of delirium, we focused on the fact that DNA methylation (DNAm) changes dynamically over the human lifespan and that epigenetic mechanisms control the expression of genes including those of cytokines. Thus, we hypothesized that epigenetic modifications specific to aging and delirium susceptibility occur in microglia; that similar modifications occur in blood; and that these epigenetic changes enhance reactions to exogenous insult, resulting in increased cytokine expression and delirium susceptibility (12). In fact, no published study has assessed DNAm and its relationship to delirium in humans, especially with genome-wide DNAm investigation.

To support this hypothesis of DNAm change along with aging among pro-inflammatory cytokines, we previously reported that DNAm levels in blood decrease along with aging on the pro-inflammatory cytokine gene *TNF-alpha*, based on 265 participants from the Grady Trauma Project (GTP) (12). We also showed that expression level of *TNF-alpha* among the same subjects increased with aging (12). This data supports our hypothesis that in pro-inflammatory cytokine genes, DNAm levels decrease along with aging, and expression level increase. In addition, using a unique dataset from our own study comparing DNAm status from neurosurgically resected live human brains followed by fluorescence-activated cell sorting (FACS), we showed that DNAm of the *TNF-alpha* universally decreases with age among the glial (neuronal negative) component, but no such patterns were found among neurons (neuronal positive component) (12). This data also supports our hypothesis that the DNAm in pro-inflammatory genes decrease in glia along with aging, making microglia potentially more prone to express those cytokine genes and to have highghtend inflammatory response when exposed to external stimuli such as surgery or infection leading to delirium.

However, what was lacking in our previous data was that we did not directly test DNAm differences between delirium cases and non-delirium controls. To fill this gap, we conducted the present study to compare DNAm status in blood from hospitalized patients with and without delirium to identify clinically useful epigenetic biomarkers for delirium from blood samples, which are routinely obtained from patients. We used blood for three reasons: 1) the function of monocytes in the blood is similar to that of microglia in that both release cytokines in response to exogenous stimulus, 2) our comparison of DNAm levels in live brain tissue (resected during neurosurgery) to those in blood from same individual collected at the same time point showed a high level of correlation genome-wide (13), and 3) our previous data showed a similar age-associated decrease in DNAm in the pro-inflammatory cytokine gene *TNF-alpha* among glia and blood (12). To accomplish these goals, we conducted the present study investigating DNAm differences in blood between delirium patients and controls without delirium.

To be comprehensive, we employed genome-wide approach using Illumina EPIC array. We first tested the pro-inflammatory cytokine gene, *TNF-alpha*, and its correlations with age between groups with and without delirium. We also conducted network analysis by using the top differentially methylated CpGs between the two groups. Lastly, we compared DNAm age between the two groups.

## METHODS AND MATERIALS

### Subjects and Sample Collection

Study participants were co-enrolled when they were enrolled for a separate, ongoing study of delirium. A more detailed overview of study participants’ recruitment process has been described previously (14). Briefly, 92 subjects were recruited for this epigenetics study between November 2017 and October 2018 at the University of Iowa Hospitals and Clinics. This study was approved by the University of Iowa’s Human Subjects Research Institutional Review Board.

### Delirium Status Definition

A more detailed overview of study participants’ phenotyping has been described previously (14). Briefly, we screened potential study participants for the presence of delirium by reviewing hospital records and by administering the Confusion Assessment Method for Intensive Care Unit (CAM-ICU) (15), the Delirium Rating Scale - Revised-98 (DRS-R-98) (16), and the Delirium Observation Screening Scale (DOSS) (17). A final decision of delirium category was conducted by a trained psychiatrist (G.S.) with detailed chart review.

### Sample Processing and Epigenetics Methods

Written informed consent was obtained, and whole blood samples were collected in EDTA tubes. All samples were stored at -80 °C. Methylome assays were performed as previously described (13). Briefly, genomic DNA was isolated from whole blood with the MasterPure™ DNA Purification kit (Epicentre, MCD85201). Genomic DNA was bisulfite-converted with the EZ DNA Methylation™ Kit (Zymo Research, D5002). DNAm of 93 samples (two samples were from one subject) was analyzed with the Infinium HumanMethylationEPIC BeadChip™ Kit (Illumina, WG-317-1002). Raw data was processed with the R packages ChAMP (18) and minfi (19, 20). One sample was filtered out because it had > 0.1 of CpG sites with detection *p*-values > 0.01, and CpG sites were filtered out as described below with ChAMP. As a result, 92 samples and 701,196 probes remained. Then, quality control and exploratory analysis were conducted. The density and multidimensional scaling plots showed 5 outliers. Two of them were the duplicate samples from one subject. The remaining 87 samples were processed again using ChAMP. The probes were filtered out if they 1) had a detection *p*-value > 0.01 (12,903 probes), 2) had less than 3 beads in at least 5% of samples per probe (8,280 probes), 3) had non-CpG probes (2,911 probes), 4) had SNP-related probes (21) (94,425 probes), 5) had multi-hit probes (22) (11 probes), and 6) were located in chromosomes X or Y (16,356 probes). Eighty-seven samples and 731,032 probes remained by filtering. Beta mixture quantile dilation (23) was used to normalize samples. Batch effect was corrected with the Combat normalization method (24) as implanted in the package SVA (25).

### Statistical Analysis

All statistical analyses were performed using R (26). Correlations between aging and DNAm levels of *TNF-alpha* at each CpGs were calculated with Pearson’s correlation analysis. The categorical data was calculated with chi-square test. DNAm age was calculated using the online DNAm Age Calculator (https://dnamage.genetics.ucla.edu/) (27) after filtering and the quality control process. Cell type proportions of CD8 T cells, CD4 T cells, natural killer cells, B cells, monocytes, and granulocytes were estimated also using the online DNAm Age Calculator (https://dnamage.genetics.ucla.edu/) (27) by using the method described in the previous study (28). Differential DNA methylation at the level of individual CpGs was analyzed by RnBeads using the limma method (29, 30). Age, gender, and cell type proportions were included as covariates. Network analysis was conducted using the R package missMethyl (31) by correcting different numbers of probes per gene on the array for Gene Ontology (GO) and Kyoto Encyclopedia of Genes and Genomes (KEGG) enrichment analysis.

## RESULTS

### Study Subjects Demographics

43 delirium case and 44 non-delirium control subjects were enrolled (age average 70.2 years, SD 10.2, range 42–101 years). Age and gender proportions were not statistically different between cases and controls. The total scores of the DRS-R-98 and DOSS were significantly higher in delirium cases than in non-delirium controls (Supplementary Table 1).

### Comparison of Delirium Cases vs Non-delirium Controls in the *TNF-alpha* Gene

Whole blood samples from 87 samples (43 delirium and 44 non-delirium, age range 42–101 years) after filtering were analyzed using the EPIC array for genome-wide DNAm analysis. There were no significant differences in CD8 T cells, natural killer cells, and B cells between delirium cases and non-delirium controls (Supplementary Table 1). Among them, first, we specifically tested correlation between age and DNAm levels at 24 CpGs in the *TNF-alpha* gene tested on the Illumina EPIC array as shown in Table 1. Delirium cases showed 3 significant CpGs after correction for multiple testing level (*p* < 0.05/24 = 0.00208), whereas non-delirium controls showed no significant CpGs after correction for multiple testing level.

**Table 1:**
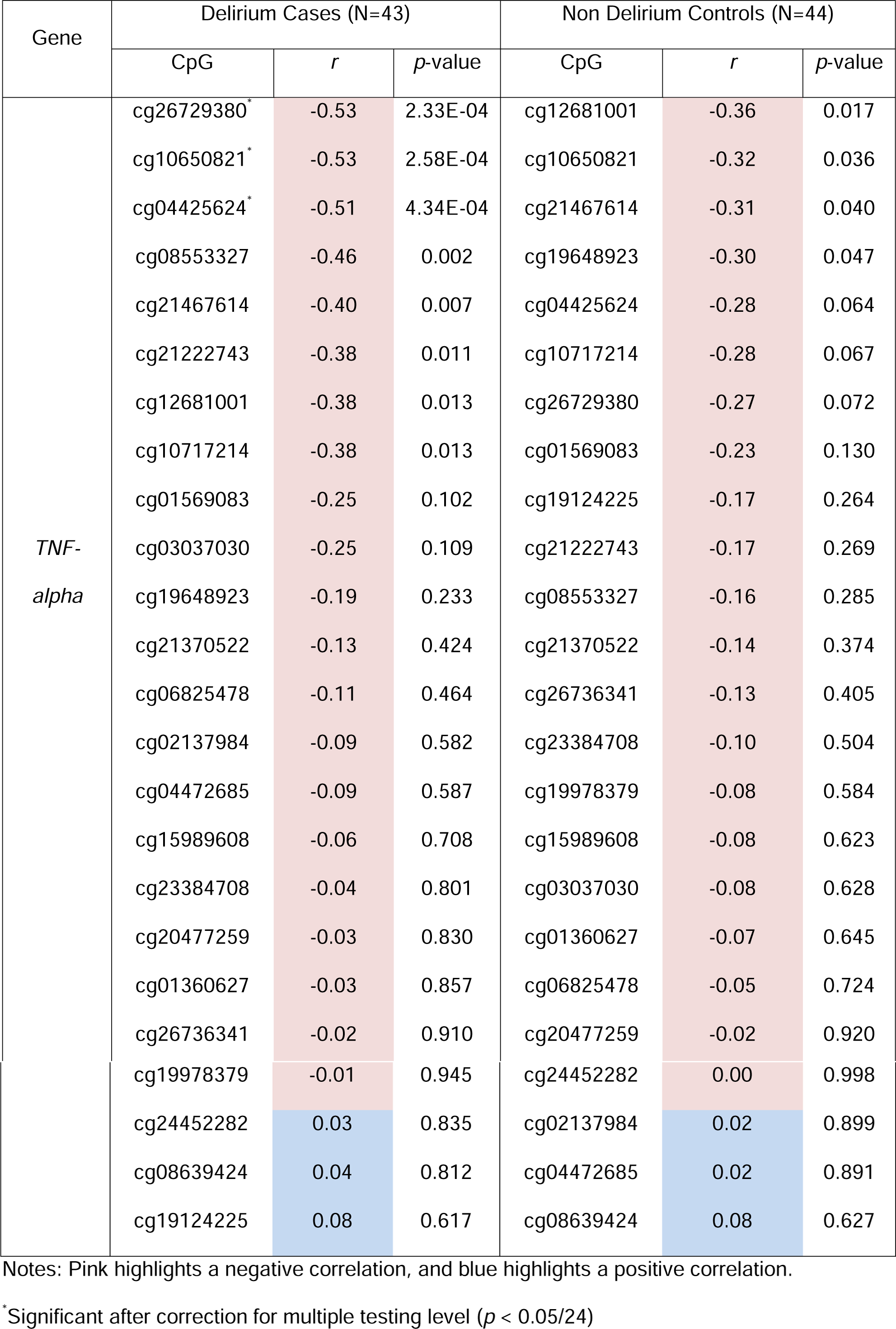
Correlation of age and blood DNAm at 24 CpGs in the *TNF-alpha* gene compared between delirium cases vs non-delirium controls

| Gene | Delirium Cases (N=43) |  |  | Non Delirium Controls (N=44) |  |  |
| --- | --- | --- | --- | --- | --- | --- |
|  | CpG | *r* | *p*-value | CpG | *r* | *p*-value |
| *TNF-alpha* | cg26729380* | -0.53 | 2.33E-04 | cg12681001 | -0.36 | 0.017 |
|  | cg10650821* | -0.53 | 2.58E-04 | cg10650821 | -0.32 | 0.036 |
|  | cg04425624* | -0.51 | 4.34E-04 | cg21467614 | -0.31 | 0.040 |
|  | cg08553327 | -0.46 | 0.002 | cg19648923 | -0.30 | 0.047 |
|  | cg21467614 | -0.40 | 0.007 | cg04425624 | -0.28 | 0.064 |
|  | cg21222743 | -0.38 | 0.011 | cg10717214 | -0.28 | 0.067 |
|  | cg12681001 | -0.38 | 0.013 | cg26729380 | -0.27 | 0.072 |
|  | cg10717214 | -0.38 | 0.013 | cg01569083 | -0.23 | 0.130 |
|  | cg01569083 | -0.25 | 0.102 | cg19124225 | -0.17 | 0.264 |
|  | cg03037030 | -0.25 | 0.109 | cg21222743 | -0.17 | 0.269 |
|  | cg19648923 | -0.19 | 0.233 | cg08553327 | -0.16 | 0.285 |
|  | cg21370522 | -0.13 | 0.424 | cg21370522 | -0.14 | 0.374 |
|  | cg06825478 | -0.11 | 0.464 | cg26736341 | -0.13 | 0.405 |
|  | cg02137984 | -0.09 | 0.582 | cg23384708 | -0.10 | 0.504 |
|  | cg04472685 | -0.09 | 0.587 | cg19978379 | -0.08 | 0.584 |
|  | cg15989608 | -0.06 | 0.708 | cg15989608 | -0.08 | 0.623 |
|  | cg23384708 | -0.04 | 0.801 | cg03037030 | -0.08 | 0.628 |
|  | cg20477259 | -0.03 | 0.830 | cg01360627 | -0.07 | 0.645 |
|  | cg01360627 | -0.03 | 0.857 | cg06825478 | -0.05 | 0.724 |
|  | cg26736341 | -0.02 | 0.910 | cg20477259 | -0.02 | 0.920 |
|  | cg19978379 | -0.01 | 0.945 | cg24452282 | 0.00 | 0.998 |
|  | cg24452282 | 0.03 | 0.835 | cg02137984 | 0.02 | 0.899 |
|  | cg08639424 | 0.04 | 0.812 | cg04472685 | 0.02 | 0.891 |
|  | cg19124225 | 0.08 | 0.617 | cg08639424 | 0.08 | 0.627 |

Notes: Pink highlights a negative correlation, and blue highlights a positive correlation.
\* Significant after correction for multiple testing level ( $p < 0.05/24$ )

### Network Analysis

Next, we directly tested genome-wide DNAm differences between delirium cases and non-delirium controls. Volcano plots for the distribution of individual CpG differences and their corresponding logarithmic transformed *p*-values are shown in Figure 1. The top 20 differentially methylated CpGs between delirium cases and non-delirium controls are shown in Supplementary Table 2. Genome-wide analysis showed a top hit at cg21295729 with genome-wide significance (*p* = 5.07E-8). This CpG is located near the gene *LDLRAD4*. Network analysis was conducted by using the CpGs with methylation level differences greater than 5.0% and *p*-value < 0.0005 (n = 753). By using those genes, network analysis showed the following results of the top 20 pathways with GO (Table 2) and KEGG analysis (Table 3). The top pathways with GO were immune response and myeloid leukocyte activation, and with KEGG were aldosterone synthesis/secretion and cholinergic synapse. Supplementary Table 3 also shows the significant pathways of GO analysis with false discovery rate (FDR)-adjusted *p*-value < 0.05.

**Table 2:**
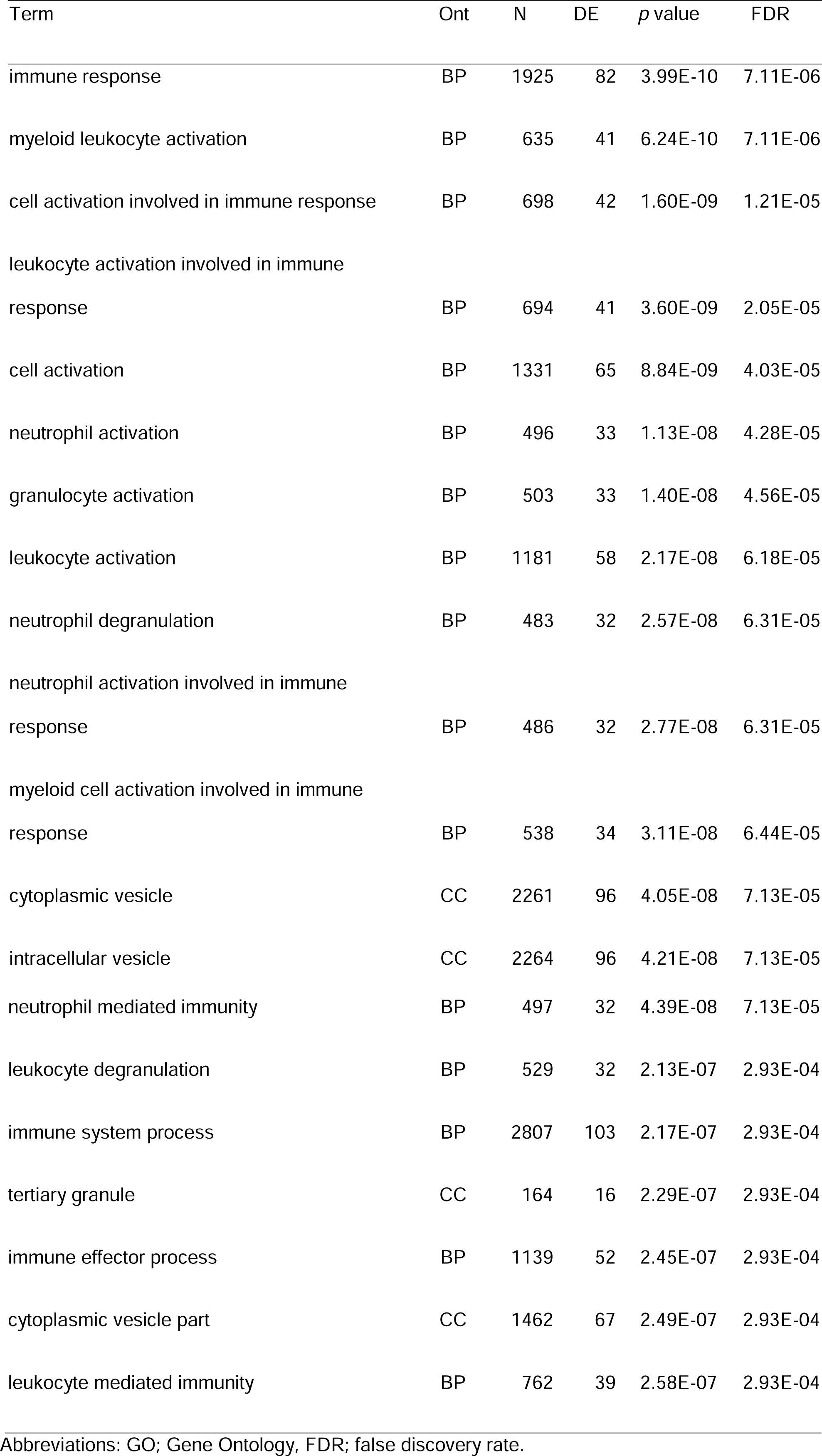
Result of the top 20 pathways of GO analysis with differentially methylated CpGs between delirium cases and non-delirium controls

**Table 3:**
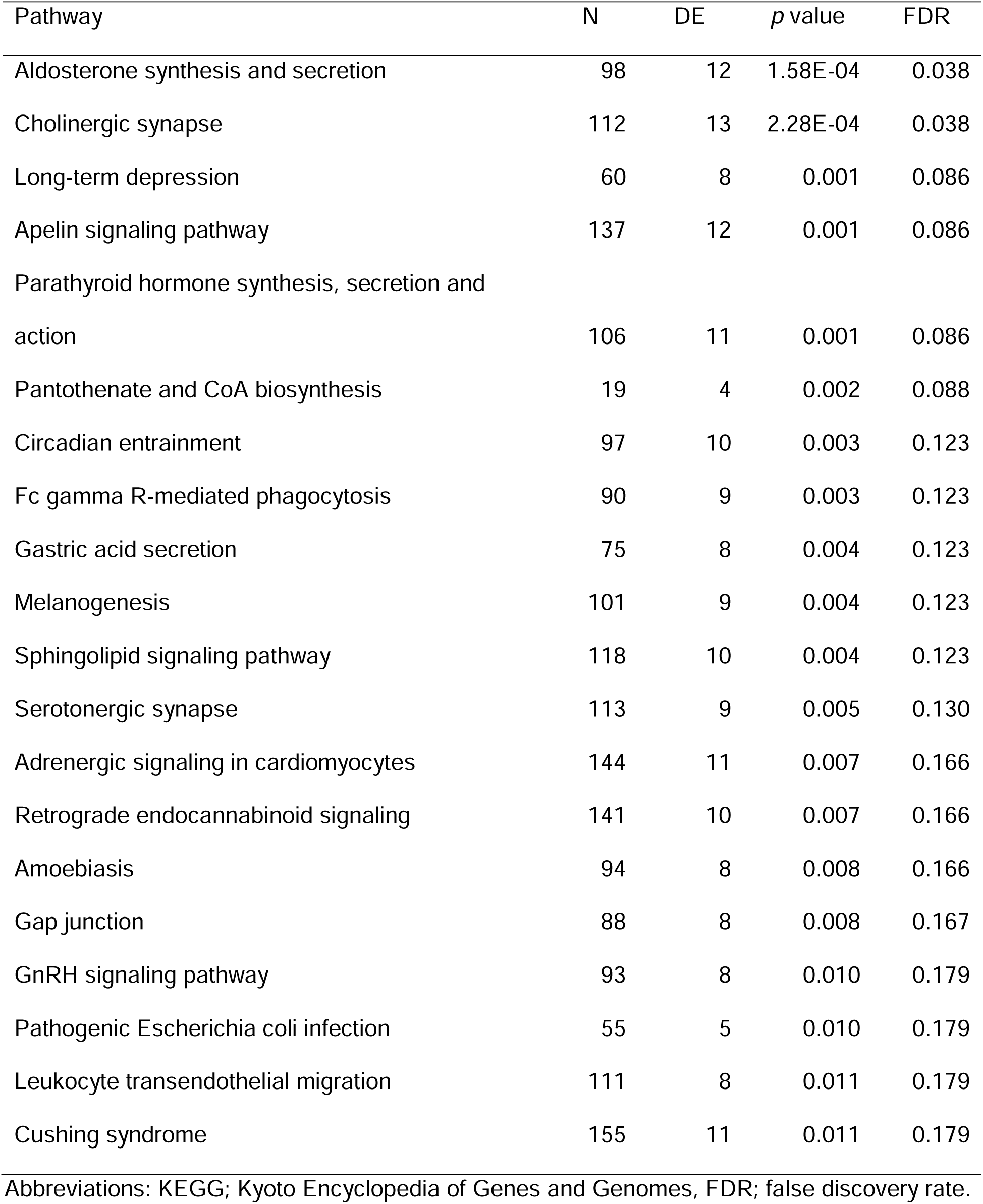
Result of the top 20 pathways of KEGG analysis with differentially methylated CpGs between delirium cases and non-delirium controls

| Pathway | N | DE | *p* value | FDR |
| --- | --- | --- | --- | --- |
| Aldosterone synthesis and secretion | 98 | 12 | 1.58E-04 | 0.038 |
| Cholinergic synapse | 112 | 13 | 2.28E-04 | 0.038 |
| Long-term depression | 60 | 8 | 0.001 | 0.086 |
| Apelin signaling pathway | 137 | 12 | 0.001 | 0.086 |
| Parathyroid hormone synthesis, secretion and action | 106 | 11 | 0.001 | 0.086 |
| Pantothenate and CoA biosynthesis | 19 | 4 | 0.002 | 0.088 |
| Circadian entrainment | 97 | 10 | 0.003 | 0.123 |
| Fc gamma R-mediated phagocytosis | 90 | 9 | 0.003 | 0.123 |
| Gastric acid secretion | 75 | 8 | 0.004 | 0.123 |
| Melanogenesis | 101 | 9 | 0.004 | 0.123 |
| Sphingolipid signaling pathway | 118 | 10 | 0.004 | 0.123 |
| Serotonergic synapse | 113 | 9 | 0.005 | 0.130 |
| Adrenergic signaling in cardiomyocytes | 144 | 11 | 0.007 | 0.166 |
| Retrograde endocannabinoid signaling | 141 | 10 | 0.007 | 0.166 |
| Amoebiasis | 94 | 8 | 0.008 | 0.166 |
| Gap junction | 88 | 8 | 0.008 | 0.167 |
| GnRH signaling pathway | 93 | 8 | 0.010 | 0.179 |
| Pathogenic Escherichia coli infection | 55 | 5 | 0.010 | 0.179 |
| Leukocyte transendothelial migration | 111 | 8 | 0.011 | 0.179 |
| Cushing syndrome | 155 | 11 | 0.011 | 0.179 |

Abbreviations: KEGG; Kyoto Encyclopedia of Genes and Genomes, FDR; false discovery rate.

**Figure 1.**
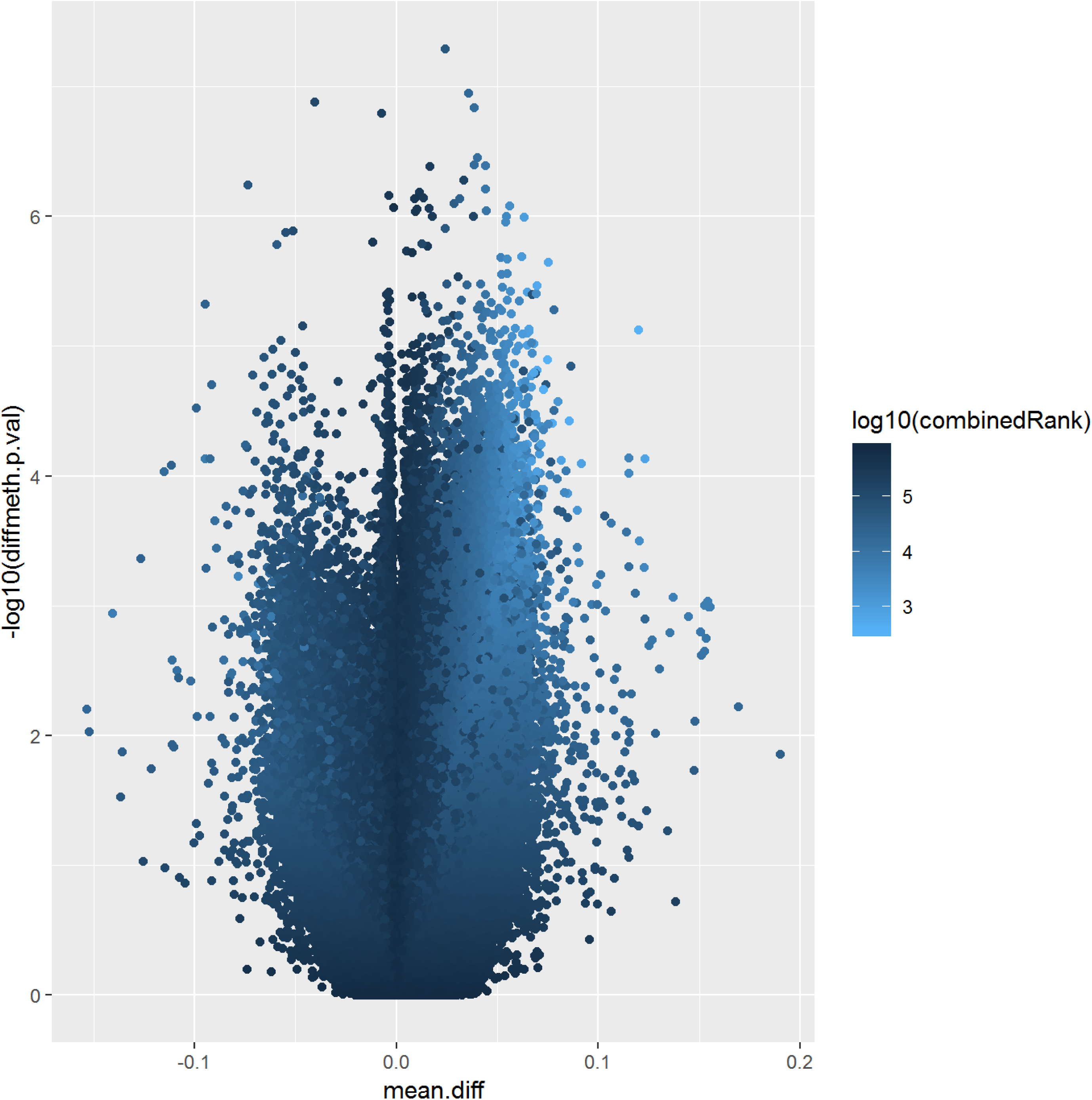
Volcano plots for the distribution of individual CpG differences and their corresponding logarithmic transformed *p*-values (n = 87)

### Comparison of Delirium Cases vs Non-delirium Controls: DNAm Age

We further tested differences of DNAm age between delirium cases and non-delirium controls. DNAm age showed signficant correlation with chronological age among delirium cases (*r* = 0.78, *p* < 0.001) and non-delirium controls (*r* = 0.67, *p* < 0.001). DNAm age among the non-delirium controls showed “slower aging” compared to the delirium cases, although there was no statistically significant difference (Figure 2).

**Figure 2.**
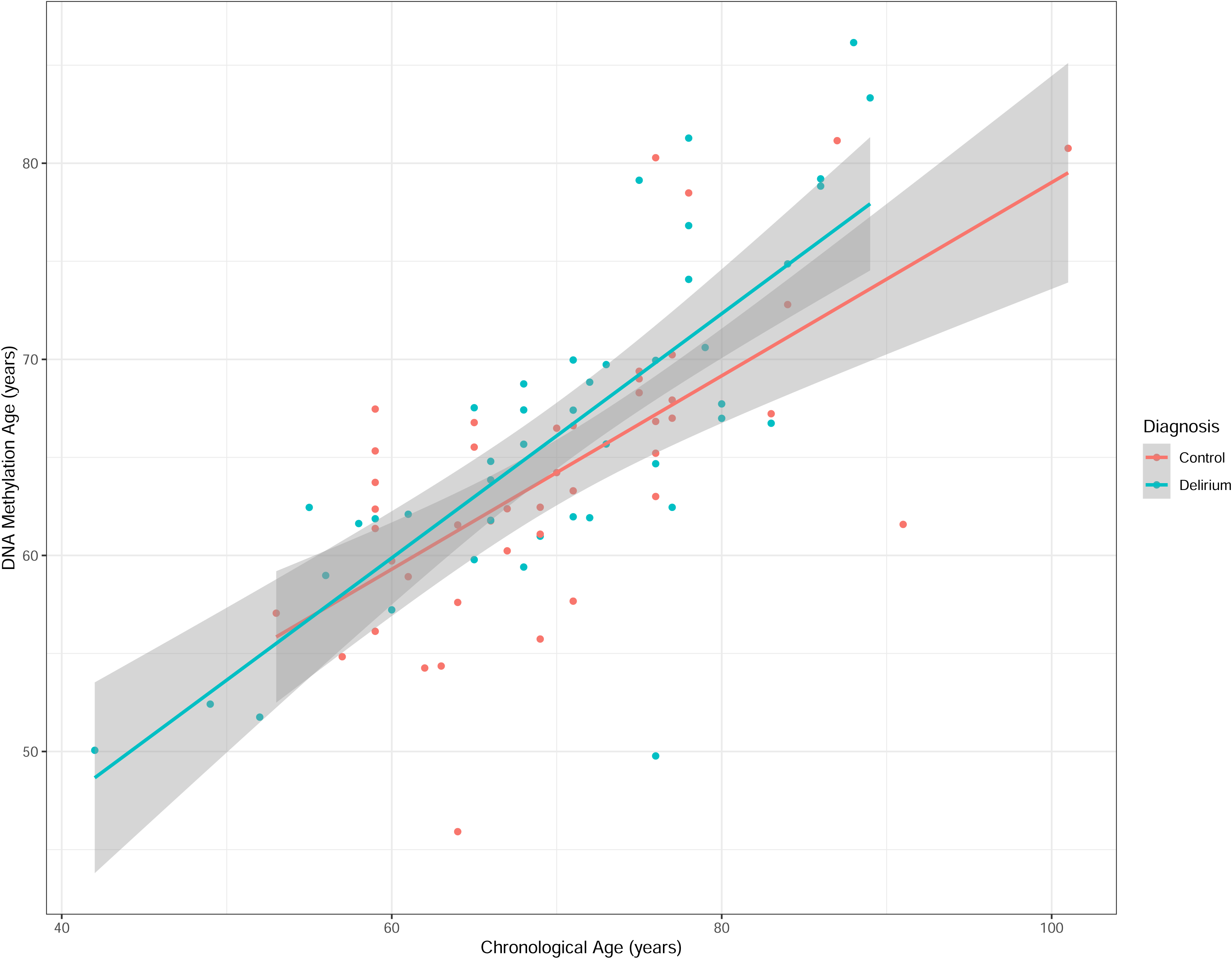
Correlation between chronological age and DNAm age in delirium (n = 43) vs controls (n = 44) Abbreviation: DNAm; DNA methylation.

## DISCUSSION

This is the first study comparing epigenetics status, especially genome-wide DNAm, between patients with and without delirium. The data presented here is consistent with our hypothesis that decreased levels of DNAm on pro-inflammatory cytokines along with aging can lead to hightened inflammation associated with delirium, whereas in patients without delirium DNAm levels remain high, and thus inflammatory reaction could be suppressed and they are protected against delirium.

In the present study, only delirium cases showed significant CpGs with multiple comparison adjusted level between age and decreasing level of *TNF-alpha* DNAm. This result is consistent with our hypothesis that the DNAm level of pro-inflammatory genes decrease along with aging in delirium patients. We speculate that delirium patients may have persistent decline of DNAm levels in the *TNF-alpha* gene along with aging, and this may have led to onset of delirium with an additional medical condition, requiring them to be admitted to a hospital. On the other hand, patients without delirium had less decline of DNAm levels in the *TNF-alpha* gene with aging, thus it might have protected them from developing delirium. To confirm this speculation, we need to conduct a large, prospective study comparing patient population as similar as possible in terms of their medication conditions. Studying surgical patients where you can collect samples before their exposure to surgery would be idealy for such investigation.

From genome-wide DNAm analysis, one genome-wide significant signal in the gene of *LDLRAD4* was identified. *LDLRAD4—*low-density lipoprotein receptor class A domain containing 4*—*functions as a negative regulator of *TGF-beta* signaling that regulates the growth, differentiation, apoptosis, motility, and matrix protein production of a lot of cell types (32). Although it was not genome-wide significant, one of the top hit CpGs was near the gene *DAPK1. DAPK1—*death associated protein kinase 1*—*functions as regulating apotosis, autophagy and inflammation (33). It is reported that *DAPK1* levels are induced by *TNF-alpha* and *interferon-gamma* (34). The other top hit CpG was near the gene *IRF8. IRF8—*interferon regulatory factor 8*—*functions as a modulating of the immune response, cell growth, and oncogenesis (35). It is reported that *IRF8* has an important role as a regulator of reactive microglia (36).

Although a potential role of these specific genes in pathophysiology of delirium requires further investigation, network analysis identified several top pathways relevant to neurofunction and inflammatory/immune processes, including immune response, leukocyte activation, neutrophil activation, and myeloid cell activation involved in immune response from GO analysis. Many pathways relevant to delirium and neural function were also identified from KEGG analysis, including cholinergic synapse, serotonergic synapse, and leukocyte transendothelial migration. The cholinergic synapse was second from the top KEGG pathways. The cholinergic system is one of the most important neurotransmitter systems in the brain, and deficiency of acetylcholine is well known to be associated with delirium (37, 38). The results of the present study are further supporting the relevance of cholinergic function in potential pathophysiology of delirium.

Furthermore, the pathways of positive regulation and regulation of interleukin-10 production, and positive regulation of interleukin-17 production, were significant (FDR-adjusted *p*-value < 0.05) from GO analysis. This result is also suggesting the role of pro-inflammatory cytokines and neuro-inflammation in pathophysiology of delirium, consistent with our hypothesis. These findings may support the validity of this epigenetic investigation of delirium pathophysiology.

We showed that DNAm age was significantly correlated with chronological age in both delirium cases and non-delirium controls. However, non-delirium cases showed relatively slower progression of DNAm aging along with chronological aging than delirium cases. DNAm age measures the accumulative effect of an epigenetic maintenance system (27) and predicts mortality (39, 40). We speculate that non-delirium controls may have a protective mechanism against DNAm aging, which can also prevent developing delirium. With a larger sample size study we can test if DNAm aging is in fact different between delirium cases versus controls.

There are several limitations in our study. First, the sample size is relatively small. To overcome this limitation, we need to increase the sample size in the future. Second, medical conditions required them to be hospitalized were diverse among study subjects including in the present study. As mentioned previously, comparing those with and without delirium after same type of surgery (post-operative delirium) would help minimize confounding factors. However, even with these limitation, the presented data showed supporting evidence of epigenetic differences in the pro-inflammatory cytokine gene *TNF-alpha* and in the immune/inflammatory response network and cholinergic system. Lastly, we used only blood samples and did not investigate brain tissues directly. However, there is a significant correlation between brain tissue and blood in DNAm levels, as shown in our previous study (13). Also, as the goal of our study is to identify potentially clinically useful biomarkers, we believe that investigating DNAm differences in blood associated with delirium is important to improve our future clinical practice.

In conclusion, to the best of our knowledge, this is the first epigenetics study of delirium. The DNAm was investigated genome-wide. The results were consistent with our previous work and hypothesis (12). Despite these limitations mentioned above, we showed evidence of epigenetic differences both at gene levels and network levels between delirium cases and non-delirium controls. This finding indicates that DNAm status in blood may become a useful epigenetic biomarker for delirium.

## ACKNOWLEDGEMENTS

This study was funded by research grants from National Science Foundation1664364 as well as the National Institute of Mental Health (K23 MH107654). The preprint version of this manuscript was posted on bioRχiv.

## REFERENCES

1. Inouye SK (2006): Delirium in older persons. N Engl J Med 354: 1157–1165.

2. Spronk PE, Riekerk B, Hofhuis J, Rommes JH (2009): Occurrence of delirium is severely underestimated in the ICU during daily care. Intensive Care Med 35: 1276–1280.

3. Pandharipande PP, Girard TD, Jackson JC, Morandi A, Thompson JL, Pun BT, et al. (2013): Long-term cognitive impairment after critical illness. N Engl J Med 369: 1306–1316.

4. McCusker J, Cole M, Abrahamowicz M, Primeau F, Belzile E (2002): Delirium predicts 12-month mortality. Arch Intern Med 162: 457–463.

5. Witlox J, Eurelings LS, de Jonghe JF, Kalisvaart KJ, Eikelenboom P, Van Gool WA (2010): Delirium in elderly patients and the risk of postdischarge mortality, institutionalization, and dementia: a meta-analysis. JAMA 304: 443–451.

6. Dillon ST, Vasunilashorn SM, Ngo L, Otu HH, Inouye SK, Jones RN, et al. (2017): Higher C-Reactive Protein Levels Predict Postoperative Delirium in Older Patients Undergoing Major Elective Surgery: A Longitudinal Nested Case-Control Study. Biol Psychiatry 81: 145–153.

7. Vasunilashorn SM, Dillon ST, Inouye SK, Ngo LH, Fong TG, Jones RN, et al. (2017): High C-Reactive Protein Predicts Delirium Incidence, Duration, and Feature Severity After Major Noncardiac Surgery. J Am Geriatr Soc 65: e109–e116.

8. Vasunilashorn SM, Ngo L, Inouye SK, Libermann TA, Jones RN, Alsop DC, et al. (2015): Cytokines and Postoperative Delirium in Older Patients Undergoing Major Elective Surgery. J Gerontol A Biol Sci Med Sci 70: 1289–1295.

9. Khan BA, Zawahiri M, Campbell NL, Boustani MA (2011): Biomarkers for delirium—a review. J Am Geriatr Soc 59: S256–S261.

10. Hovens IB, van Leeuwen BL, Nyakas C, Heineman E, van der Zee EA, Schoemaker RG (2015): Postoperative cognitive dysfunction and microglial activation in associated brain regions in old rats. Neurobiol Learn Mem 118: 74–79.

11. Xu Z, Dong Y, Wang H, Culley DJ, Marcantonio ER, Crosby G, et al. (2014): Peripheral surgical wounding and age-dependent neuroinflammation in mice. PLoS One 9: e96752.

12. Shinozaki G, Braun PR, Hing BWQ, Ratanatharathorn A, Klisares MJ, Duncan GN, et al. (2018): Epigenetics of Delirium and Aging: Potential Role of DNA Methylation Change on Cytokine Genes in Glia and Blood Along With Aging. Front Aging Neurosci 10: 311.

13. Braun PR, Han S, Hing B, Nagahama Y, Gaul LN, Heinzman JT, et al. (2019): Genomewide DNA methylation comparison between live human brain and peripheral tissues within individuals. Translational psychiatry 9: 47.

14. Shinozaki G, Chan AC, Sparr NA, Zarei K, Gaul LN, Heinzman JT, et al. (2018): Delirium detection by a novel bispectral electroencephalography device in general hospital. Psychiatry Clin Neurosci 72: 856–863.

15. Ely EW, Inouye SK, Bernard GR, Gordon S, Francis J, May L, et al. (2001): Delirium in mechanically ventilated patients: validity and reliability of the confusion assessment method for the intensive care unit (CAM-ICU). JAMA 286: 2703–2710.

16. Trzepacz PT, Mittal D, Torres R, Kanary K, Norton J, Jimerson N (2001): Validation of the Delirium Rating Scale-revised-98: comparison with the delirium rating scale and the cognitive test for delirium. J Neuropsychiatry Clin Neurosci 13: 229–242.

17. Schuurmans MJ, Shortridge-Baggett LM, Duursma SA (2003): The Delirium Observation Screening Scale: a screening instrument for delirium. Res Theory Nurs Pract 17: 31–50.

18. Morris TJ, Butcher LM, Feber A, Teschendorff AE, Chakravarthy AR, Wojdacz TK, et al. (2014): ChAMP: 450k Chip Analysis Methylation Pipeline. Bioinformatics 30: 428–430.

19. Aryee MJ, Jaffe AE, Corrada-Bravo H, Ladd-Acosta C, Feinberg AP, Hansen KD, et al. (2014): Minfi: a flexible and comprehensive Bioconductor package for the analysis of Infinium DNA methylation microarrays. Bioinformatics 30: 1363–1369.

20. Fortin JP, Triche TJ, Jr., Hansen KD (2017): Preprocessing, normalization and integration of the Illumina HumanMethylationEPIC array with minfi. Bioinformatics 33: 558–560.

21. Zhou W, Laird PW, Shen H (2017): Comprehensive characterization, annotation and innovative use of Infinium DNA methylation BeadChip probes. Nucleic Acids Res 45: e22–e22.

22. Nordlund J, Bäcklin CL, Wahlberg P, Busche S, Berglund EC, Eloranta M-L, et al. (2013): Genome-wide signatures of differential DNA methylation in pediatric acute lymphoblastic leukemia. Genome Biol 14: r105.

23. Teschendorff AE, Marabita F, Lechner M, Bartlett T, Tegner J, Gomez-Cabrero D, et al. (2013): A beta-mixture quantile normalization method for correcting probe design bias in Illumina Infinium 450 k DNA methylation data. Bioinformatics 29: 189–196.

24. Johnson WE, Li C, Rabinovic A (2007): Adjusting batch effects in microarray expression data using empirical Bayes methods. Biostatistics 8: 118–127.

25. Leek JT, Johnson WE, Parker HS, Fertig EJ, Jaffe AE, Storey JD, et al. (2019): sva: Surrogate variable analysis. R package version.

26. R Core Team (2019): R: A language and environment for statistical computing. Vienna, Austria: R Foundation for Statistical Computing.

27. Horvath S (2013): DNA methylation age of human tissues and cell types. Genome Biol 14: 3156.

28. Houseman EA, Accomando WP, Koestler DC, Christensen BC, Marsit CJ, Nelson HH, et al. (2012): DNA methylation arrays as surrogate measures of cell mixture distribution. BMC Bioinformatics 13: 86.

29. Assenov Y, Müller F, Lutsik P, Walter J, Lengauer T, Bock C (2014): Comprehensive analysis of DNA methylation data with RnBeads. Nature methods 11: 1138.

30. Ritchie ME, Phipson B, Wu D, Hu Y, Law CW, Shi W, et al. (2015): limma powers differential expression analyses for RNA-sequencing and microarray studies. Nucleic Acids Res 43: e47–e47.

31. Phipson B, Maksimovic J, Oshlack A (2016): missMethyl: an R package for analyzing data from Illumina’s HumanMethylation450 platform. Bioinformatics 32: 286–288.

32. Nakano N, Maeyama K, Sakata N, Itoh F, Akatsu R, Nakata M, et al. (2014): C18 ORF1, a novel negative regulator of transforming growth factor-β signaling. J Biol Chem 289: 12680–12692.

33. Li X, Pu J, Liu J, Chen Y, Li Y, Hou P, et al. (2018): The prognostic value of DAPK1 hypermethylation in gliomas: A site-specific analysis. Pathol Res Pract 214: 940–948.

34. Yoo HJ, Byun H-J, Kim B-R, Lee KH, Park S-Y, Rho SB (2012): DAPk1 inhibits NF-κB activation through TNF-α and INF-γ-induced apoptosis. Cell Signal 24: 1471–1477.

35. Qiu Y, Yu H, Zhu Y, Ye Z, Deng J, Su W, et al. (2017): Hypermethylation of Interferon Regulatory Factor 8 (IRF8) Confers Risk to Vogt-Koyanagi-Harada Disease. Sci Rep 7: 1007.

36. Masuda T, Tsuda M, Yoshinaga R, Tozaki-Saitoh H, Ozato K, Tamura T, et al. (2012): IRF8 is a critical transcription factor for transforming microglia into a reactive phenotype. Cell Rep 1: 334–340.

37. Hshieh TT, Fong TG, Marcantonio ER, Inouye SK (2008): Cholinergic deficiency hypothesis in delirium: a synthesis of current evidence. The Journals of Gerontology Series A: Biological Sciences and Medical Sciences 63: 764–772.

38. Maldonado JR (2013): Neuropathogenesis of delirium: review of current etiologic theories and common pathways. The American Journal of Geriatric Psychiatry 21: 1190–1222.

39. Christiansen L, Lenart A, Tan Q, Vaupel JW, Aviv A, McGue M, et al. (2016): DNA methylation age is associated with mortality in a longitudinal Danish twin study. Aging Cell 15: 149–154.

40. Marioni RE, Shah S, McRae AF, Chen BH, Colicino E, Harris SE, et al. (2015): DNA methylation age of blood predicts all-cause mortality in later life. Genome Biol 16: 25.

