## Supplementary Table 1 for "Epigenetics biomarkers of delirium: immune response, inflammatory response and cholinergic synaptic involvement evidenced by genome-wide DNA methylation analysis of delirious inpatients"

Supplementary Table 1: Demographic data and cell type proportions of delirium cases and non-delirium controls

|  | Delirium (N=43) | |  | Control (N=44) | |  | Statistical test | *p*-value |
| --- | --- | --- | --- | --- | --- | --- | --- | --- |
|  | Mean | SD |  | Mean | SD |  |  |  |
| Age (year) | 70.5 | 10.7 |  | 69.9 | 9.8 |  | *t*=0.28 | 0.777 |
| Gender (M:F) | 30:13 |  |  | 30:14 |  |  | *χ^2^*=0.00 | 1.000 |
| DNAm age (year) | 66.4 | 8.6 |  | 64.2 | 7.3 |  | *t*=1.32 | 0.189 |
| DRS-R-98 | 16.3 | 6.3 |  | 6.1 | 3.8 |  | *t*=8.70 | <0.001 |
| DOSS | 4.2 | 3.3 |  | 0.4 | 1.6 |  | *t*=6.06 | <0.001 |
| Cell type proportions |  |  |  |  |  |  |  |  |
| CD8-T | 0.04 | 0.04 |  | 0.04 | 0.04 |  | *t*=0.24 | 0.812 |
| CD4-T | 0.07 | 0.07 |  | 0.10 | 0.06 |  | *t*=2.35 | 0.021 |
| NK | 0.01 | 0.01 |  | 0.01 | 0.02 |  | *t*=0.21 | 0.830 |
| B cells | 0.01 | 0.01 |  | 0.01 | 0.02 |  | *t*=2.34 | 0.022 |
| Monocytes | 0.10 | 0.05 |  | 0.09 | 0.04 |  | *t*=0.55 | 0.587 |

Abbreviation: DNAm; DNA methylation, CD8-T; CD8 T cells, CD4-T; CD4 T cells, NK; natural killer cells, DRS-R-98; Delirium Rating Scale - Revised-98, DOSS; Delirium Observation Screening Scale.
