## Supplementary Table 2 for "Epigenetics biomarkers of delirium: immune response, inflammatory response and cholinergic synaptic involvement evidenced by genome-wide DNA methylation analysis of delirious inpatients"

Supplementary Table 2: Top 20 differentially methylated CpGs between delirium cases and non-delirium controls

| cgid | Chr | Start | Mean.  Negative | Mean.  Positive | Mean.  Diff | *p* value | Gene | Gene Group |
| --- | --- | --- | --- | --- | --- | --- | --- | --- |
| cg21295729 | chr18 | 13580042 | 0.07 | 0.04 | 0.02 | 5.07E-08 | *LDLRAD4* | Body |
| cg10518911 | chr9 | 90172046 | 0.09 | 0.05 | 0.04 | 1.13E-07 | *DAPK1* | Body |
| cg05055747 | chr12 | 94597749 | 0.83 | 0.87 | -0.04 | 1.31E-07 | *PLXNC1* | Body |
| cg21812923 | chr15 | 63127782 | 0.12 | 0.08 | 0.04 | 1.45E-07 | *TLN2* | Body |
| cg11354349 | chr12 | 53108165 | 0.03 | 0.04 | -0.01 | 1.61E-07 | *SOAT2* |  |
| cg09848074 | chr7 | 139172610 | 0.14 | 0.10 | 0.04 | 3.51E-07 | *TTC26* |  |
| cg04015794 | chr16 | 85947866 | 0.11 | 0.08 | 0.04 | 4.00E-07 | *IRF8* | Body |
| cg12400097 | chr3 | 128586320 | 0.16 | 0.12 | 0.04 | 4.05E-07 | *LOC653712* | Body |
| cg04749381 | chr12 | 11215498 | 0.97 | 0.96 | 0.02 | 4.10E-07 | *TAS2R46* | TSS1500 |
| cg00356829 | chr16 | 30025364 | 0.93 | 0.90 | 0.03 | 5.24E-07 | *FAM57B* |  |
| cg11843935 | chr8 | 128223109 | 0.76 | 0.83 | -0.07 | 5.72E-07 | *CCAT1* | Body |
| cg15412815 | chr11 | 93271088 | 0.14 | 0.09 | 0.04 | 6.12E-07 | *C11orf75* | 5'UTR |
| cg06805551 | chr3 | 196596353 | 0.94 | 0.93 | 0.01 | 6.48E-07 | *SENP5* | 5'UTR |
| cg06807989 | chr3 | 111805312 | 0.02 | 0.03 | 0.00 | 6.88E-07 | *C3orf52* | 1stExon |
| cg03231504 | chr1 | 228677993 | 0.95 | 0.93 | 0.01 | 7.18E-07 | *RNF187* | Body |
| cg00078746 | chr14 | 91700401 | 0.96 | 0.95 | 0.01 | 7.28E-07 | *GPR68* | Body |
| cg27263049 | chr8 | 145086300 | 0.10 | 0.06 | 0.03 | 7.29E-07 | *SPATC1* | TSS1500 |
| cg02010481 | chr7 | 28218524 | 0.08 | 0.05 | 0.03 | 7.95E-07 | *JAZF1* | Body |
| cg01744941 | chr4 | 79567368 | 0.17 | 0.12 | 0.06 | 8.24E-07 | *LINC01094* | Body |
| cg23623502 | chr5 | 90677000 | 0.01 | 0.01 | 0.00 | 8.47E-07 | *ARRDC3* | Body |
