## Supplementary Table 3 for "Epigenetics biomarkers of delirium: immune response, inflammatory response and cholinergic synaptic involvement evidenced by genome-wide DNA methylation analysis of delirious inpatients"

Supplementary Table 3: Result of the FDR significant pathways of GO analysis with differentially methylated CpGs between delirium cases and non-delirium controls

| Term | Ont | N | DE | *p* value | FDR |
| --- | --- | --- | --- | --- | --- |
| immune response | BP | 1925 | 82 | 3.99E-10 | 7.11E-06 |
| myeloid leukocyte activation | BP | 635 | 41 | 6.24E-10 | 7.11E-06 |
| cell activation involved in immune response | BP | 698 | 42 | 1.60E-09 | 1.21E-05 |
| leukocyte activation involved in immune response | BP | 694 | 41 | 3.60E-09 | 2.05E-05 |
| cell activation | BP | 1331 | 65 | 8.84E-09 | 4.03E-05 |
| neutrophil activation | BP | 496 | 33 | 1.13E-08 | 4.28E-05 |
| granulocyte activation | BP | 503 | 33 | 1.40E-08 | 4.56E-05 |
| leukocyte activation | BP | 1181 | 58 | 2.17E-08 | 6.18E-05 |
| neutrophil degranulation | BP | 483 | 32 | 2.57E-08 | 6.31E-05 |
| neutrophil activation involved in immune response | BP | 486 | 32 | 2.77E-08 | 6.31E-05 |
| myeloid cell activation involved in immune response | BP | 538 | 34 | 3.11E-08 | 6.44E-05 |
| cytoplasmic vesicle | CC | 2261 | 96 | 4.05E-08 | 7.13E-05 |
| intracellular vesicle | CC | 2264 | 96 | 4.21E-08 | 7.13E-05 |
| neutrophil mediated immunity | BP | 497 | 32 | 4.39E-08 | 7.13E-05 |
| leukocyte degranulation | BP | 529 | 32 | 2.13E-07 | 2.93E-04 |
| immune system process | BP | 2807 | 103 | 2.17E-07 | 2.93E-04 |
| tertiary granule | CC | 164 | 16 | 2.29E-07 | 2.93E-04 |
| immune effector process | BP | 1139 | 52 | 2.45E-07 | 2.93E-04 |
| cytoplasmic vesicle part | CC | 1462 | 67 | 2.49E-07 | 2.93E-04 |
| leukocyte mediated immunity | BP | 762 | 39 | 2.58E-07 | 2.93E-04 |
| myeloid leukocyte mediated immunity | BP | 546 | 32 | 3.46E-07 | 3.75E-04 |
| response to lipopolysaccharide | BP | 323 | 23 | 4.30E-07 | 4.45E-04 |
| tertiary granule membrane | CC | 73 | 11 | 4.51E-07 | 4.46E-04 |
| cytoplasm | CC | 11324 | 322 | 5.37E-07 | 4.94E-04 |
| secretory granule | CC | 829 | 41 | 5.42E-07 | 4.94E-04 |
| response to molecule of bacterial origin | BP | 336 | 23 | 7.23E-07 | 6.24E-04 |
| regulated exocytosis | BP | 785 | 42 | 7.40E-07 | 6.24E-04 |
| phagocytosis | BP | 236 | 22 | 1.22E-06 | 9.89E-04 |
| secretory granule membrane | CC | 294 | 22 | 1.78E-06 | 0.001 |
| intracellular signal transduction | BP | 2802 | 112 | 2.52E-06 | 0.002 |
| secretion by cell | BP | 1467 | 64 | 3.06E-06 | 0.002 |
| defense response to fungus | BP | 39 | 6 | 3.15E-06 | 0.002 |
| defense response to bacterium | BP | 243 | 14 | 4.72E-06 | 0.003 |
| response to bacterium | BP | 606 | 29 | 5.31E-06 | 0.004 |
| exocytosis | BP | 893 | 44 | 5.96E-06 | 0.004 |
| positive regulation of immune system process | BP | 978 | 45 | 7.73E-06 | 0.005 |
| secretion | BP | 1601 | 67 | 7.85E-06 | 0.005 |
| vesicle-mediated transport | BP | 1943 | 81 | 7.93E-06 | 0.005 |
| secretory vesicle | CC | 975 | 44 | 9.49E-06 | 0.006 |
| regulation of immune response | BP | 900 | 41 | 1.20E-05 | 0.007 |
| anion binding | MF | 2726 | 109 | 1.57E-05 | 0.009 |
| specific granule | CC | 160 | 14 | 2.28E-05 | 0.012 |
| leukocyte differentiation | BP | 500 | 29 | 2.51E-05 | 0.013 |
| regulation of immune system process | BP | 1416 | 57 | 2.59E-05 | 0.013 |
| antimicrobial humoral response | BP | 112 | 8 | 3.18E-05 | 0.016 |
| response to fungus | BP | 51 | 6 | 3.53E-05 | 0.017 |
| response to other organism | BP | 903 | 36 | 4.34E-05 | 0.020 |
| positive regulation of immune response | BP | 705 | 34 | 4.38E-05 | 0.020 |
| leukocyte migration involved in inflammatory response | BP | 11 | 4 | 4.41E-05 | 0.020 |
| response to external biotic stimulus | BP | 905 | 36 | 4.43E-05 | 0.020 |
| whole membrane | CC | 1624 | 68 | 5.83E-05 | 0.026 |
| specific granule membrane | CC | 91 | 10 | 6.20E-05 | 0.027 |
| cellular response to ammonium ion | BP | 63 | 9 | 6.32E-05 | 0.027 |
| inflammatory response | BP | 718 | 31 | 6.60E-05 | 0.028 |
| neutrophil aggregation | BP | 2 | 2 | 6.70E-05 | 0.028 |
| negative regulation of leukocyte apoptotic process | BP | 50 | 7 | 6.96E-05 | 0.028 |
| cytoplasmic part | CC | 9492 | 271 | 7.76E-05 | 0.031 |
| vesicle | CC | 3785 | 125 | 8.07E-05 | 0.032 |
| response to biotic stimulus | BP | 935 | 36 | 9.61E-05 | 0.037 |
| transport | BP | 4915 | 157 | 9.82E-05 | 0.037 |
| cellular response to molecule of bacterial origin | BP | 206 | 14 | 1.07E-04 | 0.040 |
| positive regulation of interleukin-10 production | BP | 30 | 5 | 1.09E-04 | 0.040 |
| positive regulation of interleukin-17 production | BP | 15 | 4 | 1.15E-04 | 0.042 |
| regulation of interleukin-10 production | BP | 45 | 6 | 1.27E-04 | 0.045 |
| chloride channel regulator activity | MF | 17 | 5 | 1.36E-04 | 0.048 |

Abbreviation: GO; Gene Ontology, FDR; false discovery rate.
